# Molecular signaling pathways underlying schizophrenia

**DOI:** 10.1101/2020.06.23.156653

**Authors:** Jari Tiihonen, Marja Koskuvi, Markku Lähteenvuo, Kalevi Trontti, Ilkka Ojansuu, Olli Vaurio, Tyrone D. Cannon, Jouko Lönnqvist, Sebastian Therman, Jaana Suvisaari, Lesley Cheng, Antti Tanskanen, Heidi Taipale, Šárka Lehtonen, Jari Koistinaho

## Abstract

The molecular pathophysiological mechanisms underlying schizophrenia have remained unknown, and no treatment exists for primary prevention. Studies using stem cell-derived neurons have investigated differentially expressed genes (DEGs) and GO and KEGG pathways between patients and controls, but not analyzed data-driven causal molecular pathways involved. We used Ingenuity Pathway Analysis (IPA) to analyze canonical and causal pathways in two different datasets, including patients from Finland and USA. The most significant findings in canonical pathway analysis were observed for glutamate receptor signaling, hepatic fibrosis, and glycoprotein 6 (GP6) pathways in the Finnish dataset, and GP6 and hepatic fibrosis pathways in the US dataset. In data-driven causal pathways, *ADCYAP1, ADAMTS*, and *CACNA* genes were involved in the majority of the top 10 pathways differentiating patients and controls in both Finnish and US datasets. In contrast, no dopamine-specific genes were consistently involved. Results from a Finnish nation-wide database showed that the risk of schizophrenia relapse was 41% lower among first-episode patients during the use of losartan, the master regulator of an *ADCYAP1, ADAMTS*, and *CACNA* -related pathway, compared to those time periods when the same individual did not use the drug. This association was not attributable to general adherence to drug treatments. The results from the two independent datasets suggest that the GP6 signaling pathway, and the *ADCYAP1, ADAMTS*, and *CACNA* -related purine, oxidative stress, and glutamatergic signaling pathways are primary pathophysiological alterations in schizophrenia among patients with European ancestry. While no reproducible dopaminergic alterations were observed, the results imply that agents such as losartan, and ADCYAP1/PACAP -deficit alleviators, such as metabotropic glutamate 2/3 agonist MGS0028 and 5-HT7 antagonists – which have shown beneficial effects in an experimental *Adcyap1^-/-^* mouse model for schizophrenia – could be potential treatments before the full manifestation of illness involving dopaminergic abnormalities.

## Introduction

The first effective pharmacological treatments for schizophrenia were discovered more than 60 years ago, and ever since, all of them have been full or partial dopamine D_2_-receptor antagonists. Therefore, on the basis of the dopamine hypothesis, it was believed for decades that the main pathophysiology underlying the clinical phenotype was a defect in the dopaminergic system [1]. However, only a minority of patients reach full recovery or remission when treated with medications currently available, and, especially, negative and cognitive symptoms are commonly resistant to existing antipsychotic treatments [1]. There is increasing evidence that, in addition to dopaminergic defects, also abnormal glutamatergic, GABAergic, and serotoninergic signaling, as well as inflammation and oxidative stress contribute to the pathophysiology of schizophrenia [1,2].

Although large-scale genetic studies have been able to reveal hundreds of gene polymorphisms associated with schizophrenia [3], the attributable fraction of each gene is only a few per mille at best, and the findings have not been able to elucidate how these variants contribute to complex molecular pathways in abnormal brain development leading to mental illness. The same applies to postmortem studies [4–6] since postmortem brain tissue is affected by treatment and course of the illness. However, the pathophysiology of neurological and psychiatric disorders can be modeled using human induced pluripotent stem cell (hiPSC)-derived neurons. So far, these studies [7,8] have tried to identify abnormalities related to illness by analyzing how differentially expressed genes (DEGs) interact in established pathways such as GO and KEGG, based on previous literature on gene functions, but no reproducible results identifying novel potential treatments have been published using these analyses. Ingenuity Pathway Analysis (IPA) is a data-driven method that may increase the possibilities to detect disease-specific abnormalities [9]. We used IPA in two independent hiPSC-neuron datasets as well as a national pharmacoepidemiological database in order to find potential novel treatments based on molecular pathways underlying schizophrenia, and to study whether any of these findings are consistent and reproducible.

## Material and Methods

### Pathway analysis

We applied knowledge-based Ingenuity Pathway Analysis [9] to investigate two published DEG datasets comparing hiPSC-derived neurons from patients with schizophrenia to those of unrelated healthy individuals [8,10]. In addition, we aimed to include a comparison of monozygotic twin pairs discordant for schizophrenia, but this comparison yielded only one DEG surviving correction for multiple comparisons in ref [8], and no pathway analysis was possible. In order to have a larger number of DEGs for pathway analysis, we made a new comparison with neuronal cells dissociated for final maturation from neuroprogenitors maintained in culture for 12 weeks, including the same five twin pairs as earlier (in the previous studies, hiPSC-derived neuronal cells were maintained in culture for 6 [10] and 10 [8] weeks). In total, these datasets included results from 37 individuals (16 patients, 16 healthy unrelated controls, and 5 unaffected monozygotic twins). All 15 individuals included in the Finnish dataset were Caucasian and of Finnish origin. In the US dataset, 9 of 11 patients were Caucasian and 2 Caucasian-Hispanic, and 5 of 11 controls were Caucasian, the rest 6 having mixed ethnicity (Supplementary data 1 [10]). From all three datasets, genes with a minimum of 2-fold change in expression and adjusted *p*<0.05 (232 DEGs/231 genes mapped by IPA [8] for affected twin vs. controls, and 356 DEGs/352 mapped by IPA for unaffected twin vs. controls; new hiPSC-derived neuron data made for this work, 69 DEGs/all mapped by IPA) or nominal *p*<0.05 (108 DEGs/100 mapped by IPA [10]) were used as input for IPA core analysis, and run with default settings against data content version 49932394 (dated 2019-11-15). In causal pathway analysis, the software used previous knowledge on molecular pathways to identify a master regulator, which regulated a cluster of genes via several upstream regulators.

DEG lists applied to IPA, and the newly produced RNA-sequencing data are stored in University of Helsinki data cloud (https://datacloud.helsinki.fi), accessible via the link: https://datacloud.helsinki.fi/index.php/s/MjdMNcW2QMt7zFk Hoffman et al. [10] SZ vs control DEG data of neurons was downloaded from https://www.nature.com/articles/s41467-017-02330-5#Sec35 (Supplementary Data 8).

### hiPSC cell culture work of monozygotic twins

#### hiPSC cell culture and neural differentiation

Methods of cell culturing, RNA sequencing, and DEG analysis followed those published in ref. [8]. Briefly, hiPSCs cultured in E8 medium (Thermo Fisher Scientific) on Matrigel–coated dishes were differentiated to neurons using dual SMAD inhibitors SB431542 and LDN193189 (Selleckchem) for 10 days. The neuroprogenitor cells formed the spheres and were maintained in floating culture for 12 weeks instead of 10, as previously published [8]. The cells were dissociated with Accutase for the final maturation in monolayer culture for one week before RNA collection.

#### RNA isolation and transcriptomic analysis

RNA was isolated from the plated neurons by RNeasy mini kit (Qiagen) according to the manufacturer’s protocol. The Agilent 2100 Bioanalyzer confirmed the required quality of RNA before the whole transcriptome sequencing was performed on the Illumina Hiseq 2500 by the Sequencing laboratory of Institute for Molecular Medicine Finland FIMM Technology Centre, University of Helsinki. The sequencing data normalization and DEG analysis were performed using R package DESeq2, v. 1.16.1, as previously.

### Pharmacoepidemiological analysis

The details of the method are shown in Supplementary Methods and Supplementary Tables 1, 2, and 3.

## Results

Table 1a shows the top 10 canonical pathways in the Finnish dataset [8] comparing patients and unrelated healthy controls, and all pathways are shown in Supplementary Table 4. The most significant pathways were glutamate receptor signaling and white adipose tissue browning pathway (p = 1.3 x 10^-4^ in both), the first involving glutamate ionotropic receptor kainate type subunit 2 *(GRIK2)*, glutamate receptor ionotropic NMDA2B-subunit *(GRIN2)*, metabotropic glutamate receptor 1 (*GRM1*), vesicular glutamate transporter 1 *(SLC17A6)*, and vesicular glutamate transporter 2 *(SCL17A7)* genes. The combined effect of these altered gene expressions in the glutamate signaling pathway leads to increased postsynaptic neurotoxicity, synaptic plasticity, excitatory potentials, and receptor, as shown in Figure 1a. In the comparison between affected vs. unaffected twins in the novel Finnish dataset (prepared for this study), the most statistically significant canonical pathways were hepatic fibrosis/hepatic stellate cell activation (p = 7.9 x 10^-20^), and glycoprotein 6 (GP6) signaling (p = 1.3 x 10^-12^) pathways (Table 1b, Supplementary Table 5, Figure 1b). In the US dataset [10], the most significant finding was observed for glycoprotein 6 (GP6, p = 3.0 x 10^-6^) and hepatic fibrosis/hepatic stellate cell activation (p = 3.8 x 10^-5^) pathways, including many same genes (Table 1c, Supplementary Table 6, Figure 1c).

**Figure 1.**
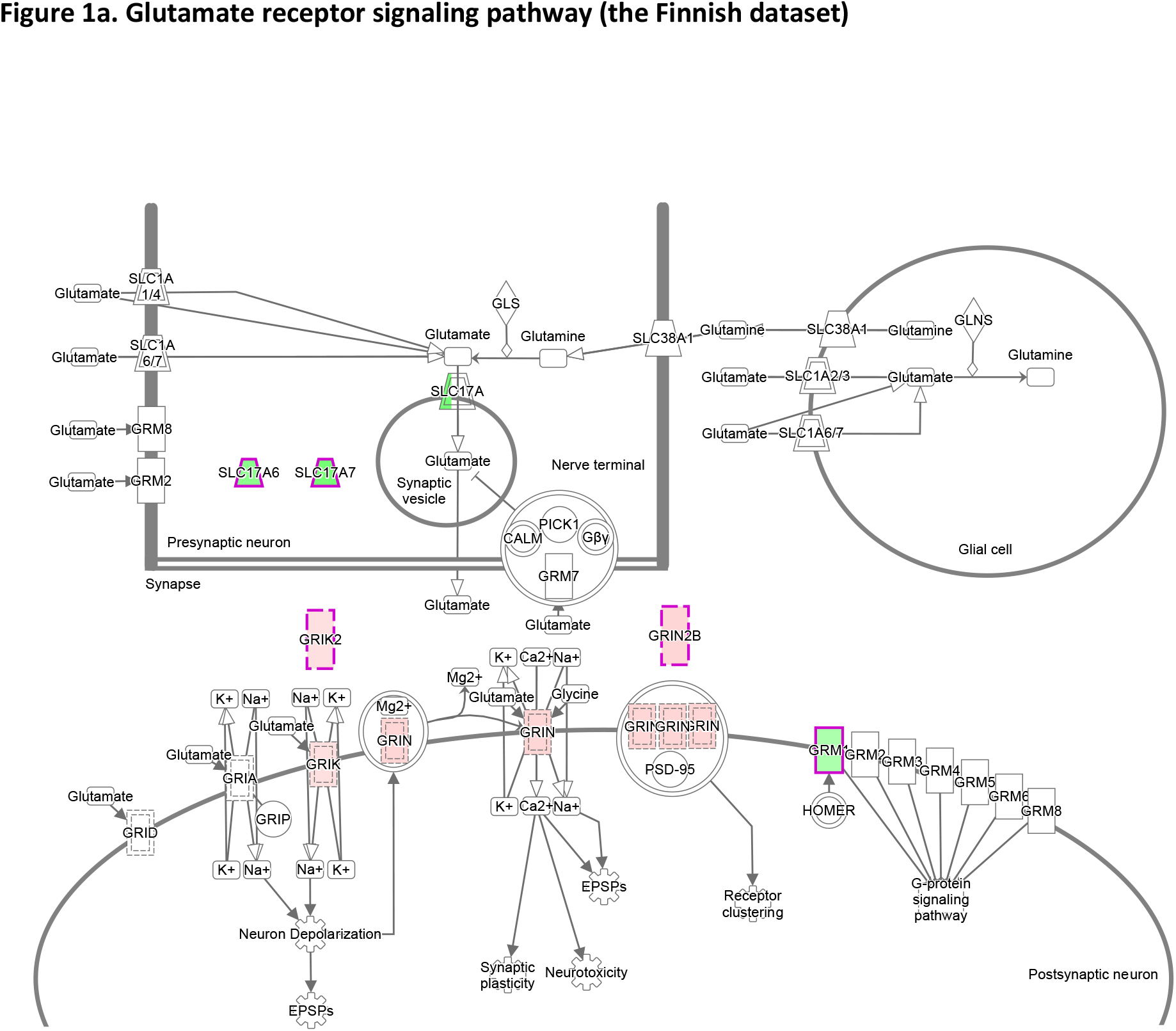
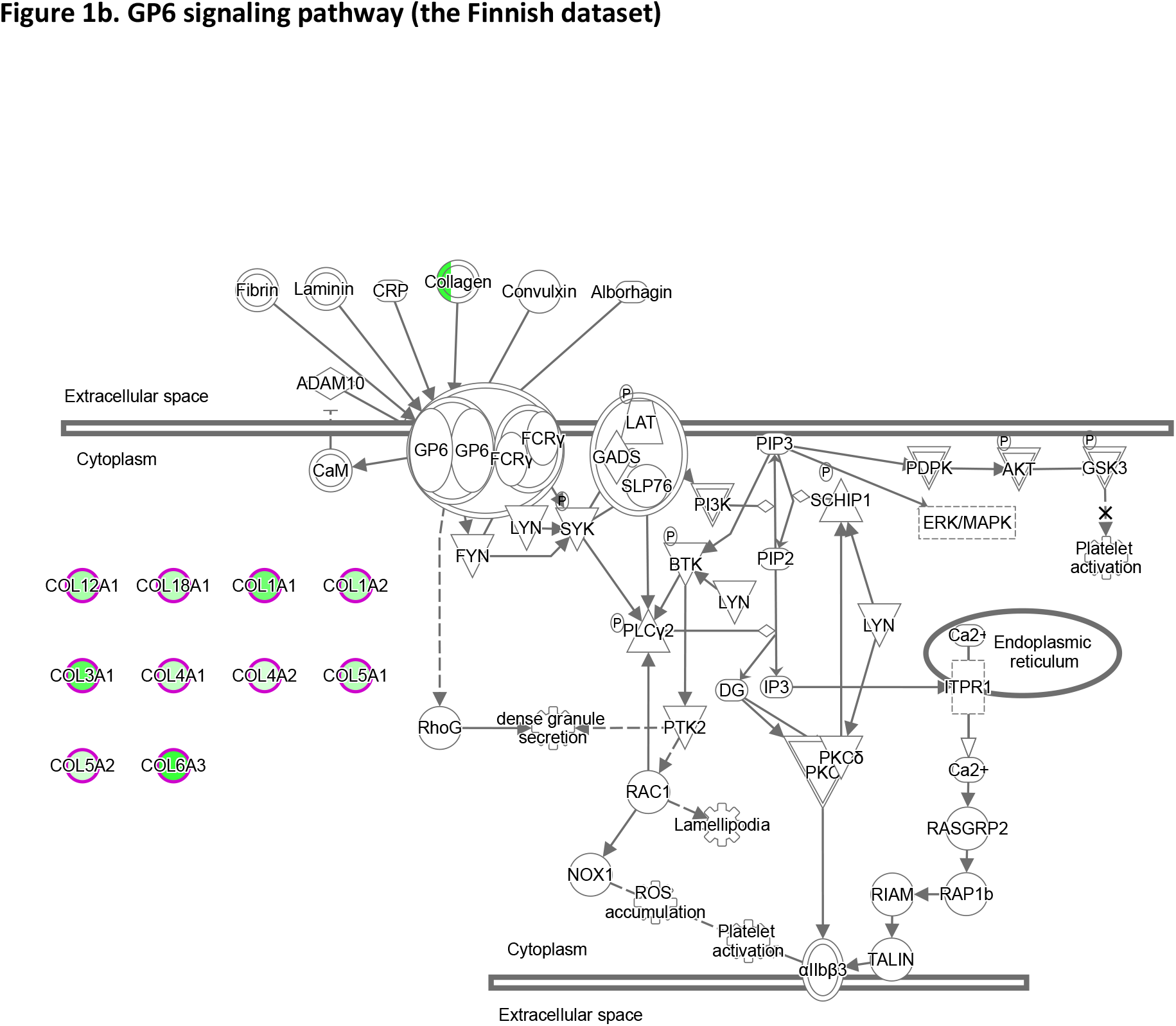
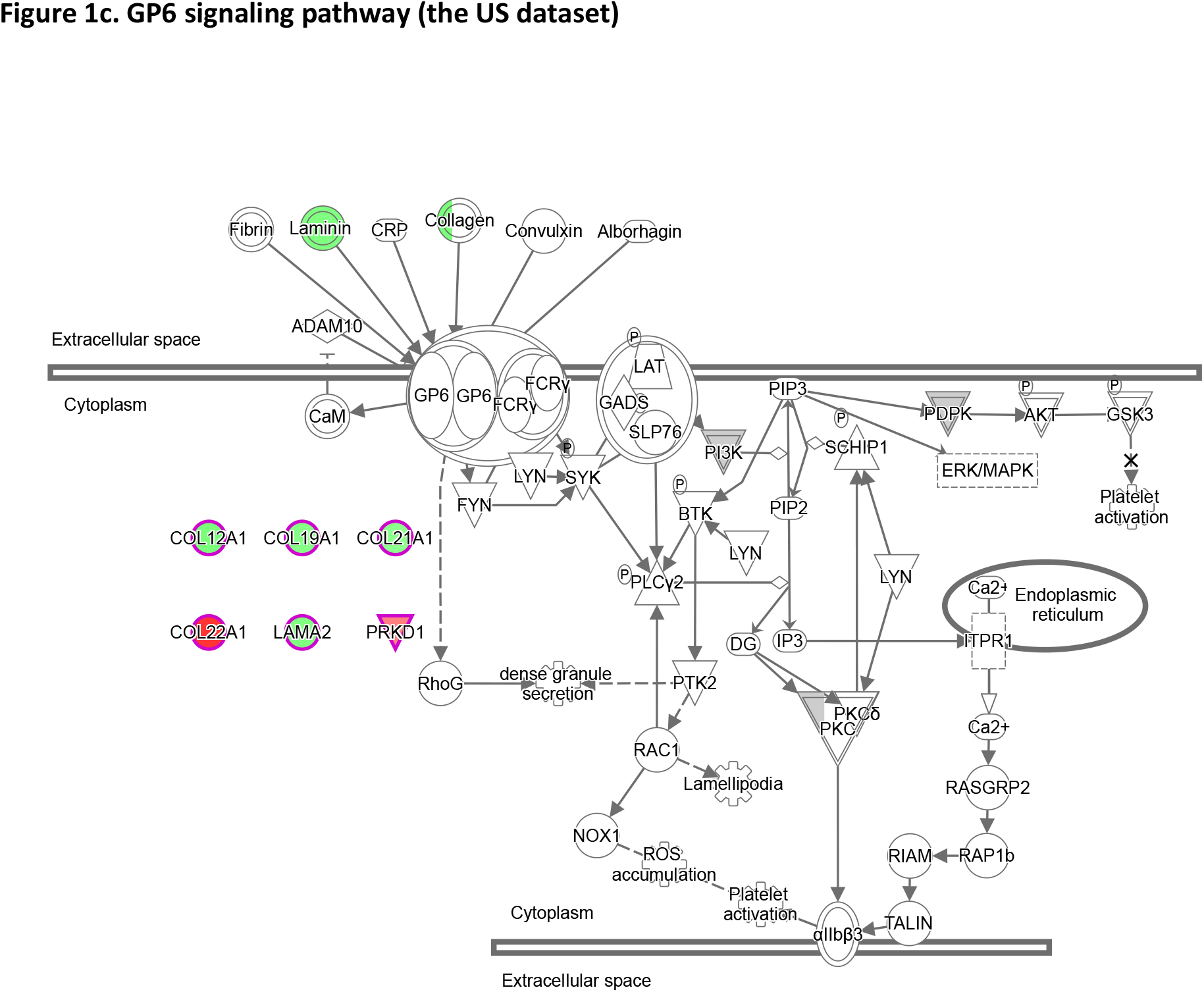
IPA canonical pathways with participating differentially expressed genes, highlighted with a violet border. Red or green color indicates that the gene or pathway member is upregulated or downregulated, respectively, in affected compared to healthy individuals. (a) Glutamate receptor signaling in the comparison of affected twins vs. unrelated healthy controls from the Finnish dataset [8], (b) GP6 signaling in comparison of affected twins vs. unaffected twins in hiPSC-neuron data created for this work, and (c) GP6 signaling from the US dataset [10] showing the comparison of patients with schizophrenia vs. healthy individuals.

**Table 1a.**
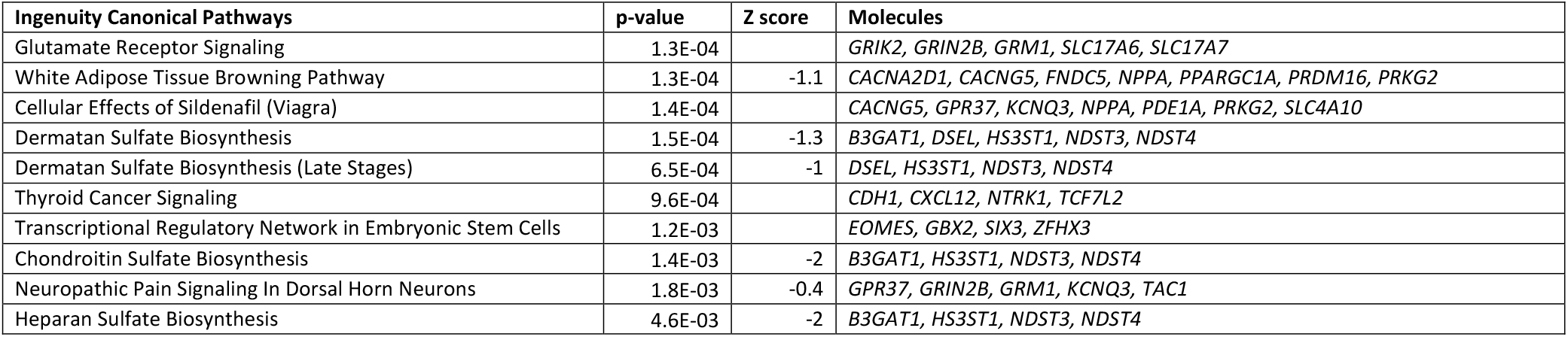
The top 10 canonical pathways observed in the comparison between patients with schizophrenia and healthy unrelated controls in the Finnish dataset [8]

**Table 1b.**
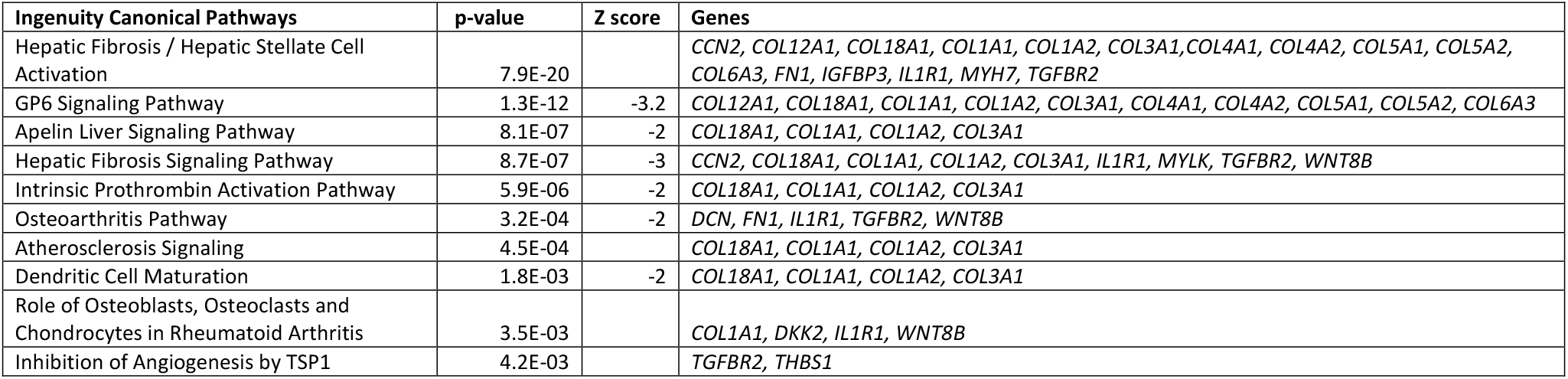
The top 10 canonical pathways in the comparison between affected and unaffected twins in the Finnish dataset [8]

**Table 1c.** The top 10 canonical pathways observed in the comparison between patients with schizophrenia and healthy unrelated controls in the US dataset [10]

| **Ingenuity Canonical Pathways** | **p-value** | **Z score** | **Genes** |
| --- | --- | --- | --- |
| GP6 Signaling Pathway | 3.0E-06 | -0.8 | *COL12A1, COL19A1, COL21A1, COL22A1, LAMA2, PRKD1* |
| Hepatic Fibrosis / Hepatic Stellate Cell Activation | 3.8E-05 |  | *COL12A1, COL19A1, COL21A1, COL22A1, LHX2, MYH14* |
| Transcriptional Regulatory Network in Embryonic Stem Cells | 1.4E-02 |  | *NEUROG1, OTX1* |
| Retinoic acid Mediated Apoptosis Signaling | 1.7E-02 |  | *CFLAR, RXRG* |
| Cardiac Hypertrophy Signaling (Enhanced) | 2.3E-02 | 1 | *ADRA1A, CACNA1A, FZD8, MPPED2, PRKD1* |
| ILK Signaling | 2.6E-02 |  | *DSP, FLNC, MYH14* |
| Superoxide Radicals Degradation | 2.6E-02 |  | *CAT* |
| FcγRIIB Signaling in B Lymphocytes | 2.6E-02 |  | *CACNA1A, INPP5D* |
| VDR/RXR Activation | 2.8E-02 |  | *PRKD1, RXRG* |
| IL-3 Signaling | 2.9E-02 |  | *INPP5D, PRKD1* |

Table 2 shows the results for the top 10 data-driven causal pathways, highlighting schizophrenia-related genes, and all pathways and genes are illustrated in Supplementary Table 4 (Finnish dataset) and Supplementary Table 6 (US dataset). Both datasets included a large number of genes linked previously with schizophrenia (see Supplementary Table 7). In the Finnish dataset, the most significant pathways were regulated by P2RY11 purine receptor and losartan potassium (angiotensin II receptor antagonist used as a treatment for hypertonia), both including several genes coding glutamate receptors and transporters (Supplementary Figures 1 and 2). In the US dataset, the strongest findings were observed for pathways regulated by cytosolic phospholipase Cpla2 and the alpha-adrenergic receptor (Supplementary Figures 3 and 4). *ADCYAP1, ADAMTS*, and *CACNA* genes were included in most of the top 10 pathways in both Finnish and US datasets. Still, dopamine-related genes *(DRD2* and *COMT)* were observed only in the Finnish dataset, and GABA-related genes *(GABRA2)* only in the US dataset (Table 2, Figure 2). Collagen genes were included in numerous pathways in both datasets, implying abnormal cell adhesion and extracellular matrix in schizophrenia [11]. Figure 2 illustrates the genes related to purine receptor, oxidative stress, and glutamate receptor signaling.

**Figure 2.**
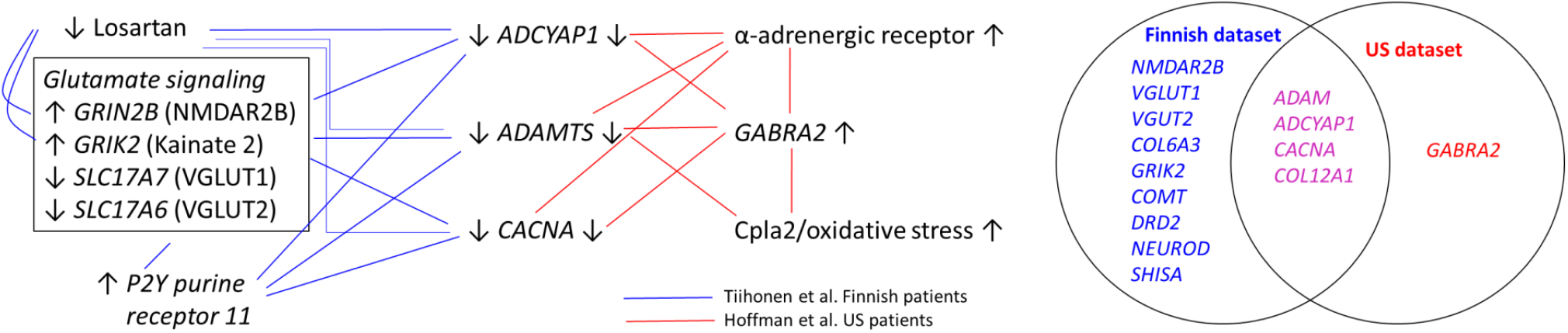
The network of differentially expressed genes in the Finnish and US datasets. Arrows indicate increased (up) and decreased (down) activity in the comparison between patients with schizophrenia and healthy controls. For example, higher expression of 2PY purine receptor 11 and x-adrenergic receptors, and lower losartan-type activity in patients is associated with abnormal gene expressions. The Venn diagram shows the visualization of overlap of altered gene expression related to schizophrenia in the Finnish (blue) and US (red) datasets. Common genes for both datasets are indicated by violet color.

**Table 2.** The top 10 causal pathways in the Finnish [8] and US [10] datasets comparing patients and healthy unrelated controls. Common genes/members of a gene family in both datasets are in bold.

| Tiihonen et al. dataset [8] |  |  |  |  | Hoffman et al. dataset [10] |  |  |  |  |
| --- | --- | --- | --- | --- | --- | --- | --- | --- | --- |
| Master Regulator | Molecule Type | p-value of overlap | Network bias-corrected p-value | Target molecules in dataset | Master Regulator | Molecule Type | p-value of overlap | Network bias-corrected p-value | Target molecules in dataset |
| P2RY11 | G-protein coupled receptor | 3.9E-10 | 0.0001 | **ADAMTS9, ADCYAP1, CACNA2D1, COMT, GRIK2, GRIN2B, NEUROD1, NEUROD6, SHISA6, SLC17A6, SLC17A7** | Cpla2 | group | 2.9E-08 | 0.0003 | **ADAMTS2, GABRA2** |
| losartan potassium | chemical drug | 8.6E-10 | 0.0001 | **ADAMTS9, ADCYAP1, CACNA2D1, COMT, GRIK2, GRIN2B, NEUROD1, NEUROD6, SLC17A6** | alpha-adrenergic receptor | group | 5.7E-08 | 0.0004 | **ADAMTS2, ADCYAP1, CACNA1A** |
| TAF4 | transcription regulator | 9.8E-10 | 0.0001 | **ADCYAP1, CACNA2D1, COMT, DRD2, GRIK2, NEUROD1, NEUROD6, SHISA6, SLC17A6** | GCKR | other | 8.8E-08 | 0.0003 | **ADAMTS2, ADCYAP1, CACNA1A** |
| PRKD2 | kinase | 1.6E-09 | 0.0001 | **ADAMTS9, ADCYAP1, CACNA2D1, COMT, GRIK2, NEUROD1, NEUROD6, SLC17A6** | costunolide | chemical - endogenous non-mammalian | 4.1E-07 | 0.0017 | **ADAMTS2, ADCYAP1, CACNA1A, GABRA2** |
| AKT1S1 | other | 1.7E-09 | 0.0001 | **ADAMTS9, ADCYAP1, CACNA2D1, COMT, NEUROD6, SLC17A6, SLC17A7** | ICL1-9 | chemical reagent | 6.7E-07 | 0.0011 | **ADAMTS2, ADCYAP1, CACNA1A** |
| CABIN1 | other | 2.8E-09 | 0.0001 | **ADAMTS9, ADCYAP1, CACNA2D1, COMT, DRD2, GRIK2, GRIN2B, NEUROD6, SLC17A6** | Fcgr3 | group | 8.9E-07 | 0.0013 | **ADAMTS2, ADCYAP1, CACNA1A, GABRA2** |
| CaMK I | group | 7.2E-09 | 0.0002 | **ADAMTS9, ADCYAP1, CACNA2D1, COMT, DRD2, GRIK2, GRIN2B, NEUROD1, NEUROD6, SLC17A6** | NCSTN | peptidase | 1.1E-06 | 0.0024 | **ADAMTS2, ADCYAP1, CACNA1A** |
| Retinoic acid-RAR-RXR | complex | 1.1E-08 | 0.0001 | **ADAMTS9, CACNA2D1, COMT, DRD2, GRIK2, NEUROD1, NEUROD6, SHISA6, SLC17A6** | L-tyrosine | chemical - endogenous mammalian | 1.2E-06 | 0.0007 | **ADAMTS2, CACNA1A** |
| GEM | enzyme | 1.2E-08 | 0.0001 | **CACNA2D1, GRIK2, NEUROD1, NEUROD6, SLC17A6** | astaxanthin | chemical drug | 1.9E-06 | 0.0007 | **CACNA1A** |
| RP 73401 | chemical toxicant | 1.5E-08 | 0.0004 | **ADAMTS9, ADCYAP1, CACNA2D1, COMT, GRIK2, GRIN2B, NEUROD6, SLC17A6** | ADRB1 | G-protein coupled receptor | 1.9E-06 | 0.0009 | **ADCYAP1, CACNA1A** |

In the comparison between Finnish affected vs. unaffected twins, the most significant pathway was complement C3/4 receptor-like 1 (CR1L) regulated pathway (p = 2.6 x 10^-18^, Supplementary Table 5, Supplementary Figure 5). There was considerable overlap of regulated genes in CR1L-regulated pathway and GP6 and hepatic fibrosis pathways since the majority of GP6 and CR1L genes were included in hepatic fibrosis/hepatic stellate cell activation pathway. Most of those common genes coded for collagen. As can be seen in Tables 1b and 1c, the most significant findings were seen for the same canonical pathways (GP6 and hepatic fibrosis/hepatic stellate cell activation) in the comparison between patients vs. controls in the US dataset, and in the comparison between affected vs. unaffected twins in the Finnish dataset.

Supplementary Table 8 shows canonical and causal pathways for comparison between unaffected twins versus healthy controls. As expected, the effect sizes were smaller than in the comparison between affected twins versus healthy controls. In the canonical pathway analysis, the comparison between affected versus unaffected twins showed the strongest finding for hepatic fibrosis and GP 6 pathways, whereas in the comparison between affected twins versus unrelated healthy controls, the most significant finding was glutamatergic signaling pathway. This is apparently explained by the fact that unaffected twins share partially the same abnormalities as their affected twins – for example, *GRIN2B* was upregulated and *CACNA2D* downregulated among them when compared with healthy controls. Therefore, the glutamatergic pathway does not show up in the comparison between affected versus unaffected twins. This suggests that glutamatergic abnormalities are associated with the shared familiar risk among affected and unaffected twins, and GP 6 pathway is associated with the actual clinical illness.

Figure 3 shows the risk of rehospitalizations due to psychosis (indicator for severe relapse) in a Finnish nationwide cohort of patients with schizophrenia. Use of any angiotensin II receptor antagonist was associated with 27% lower risk, and use of losartan with 27% (95% CI 20%–34%) lower risk of rehospitalization due to psychosis compared with those time periods in the total cohort when the same individual was not using the drug (within-individual analysis). In the incident cohort of 8,342 first-episode patients with a median age of 36 years, the corresponding decrease for losartan use was 41% (95% CI 14%–60%). The results for thiazide diuretics, used for the same somatic indication (hypertonia), indicated that the findings were not attributable to general adherence to drug treatments. Antipsychotic use was associated with only slightly lower (46% vs. 41%) risk of rehospitalization than losartan use among the incident cohort, but no beneficial outcome was observed for benzodiazepine use. The raw data are shown in Supplementary Tables 9a and 9b.

**Figure 3.**
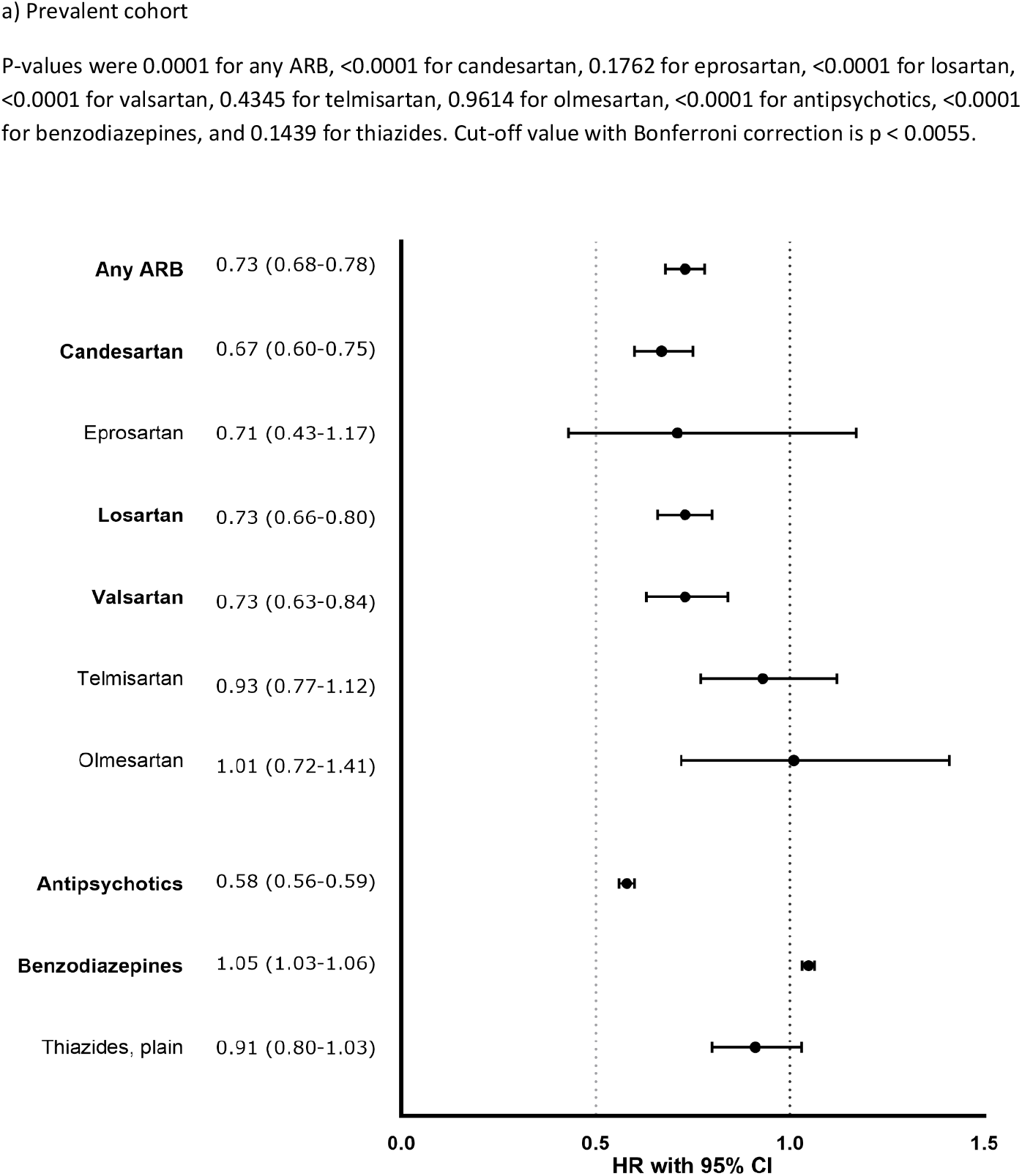
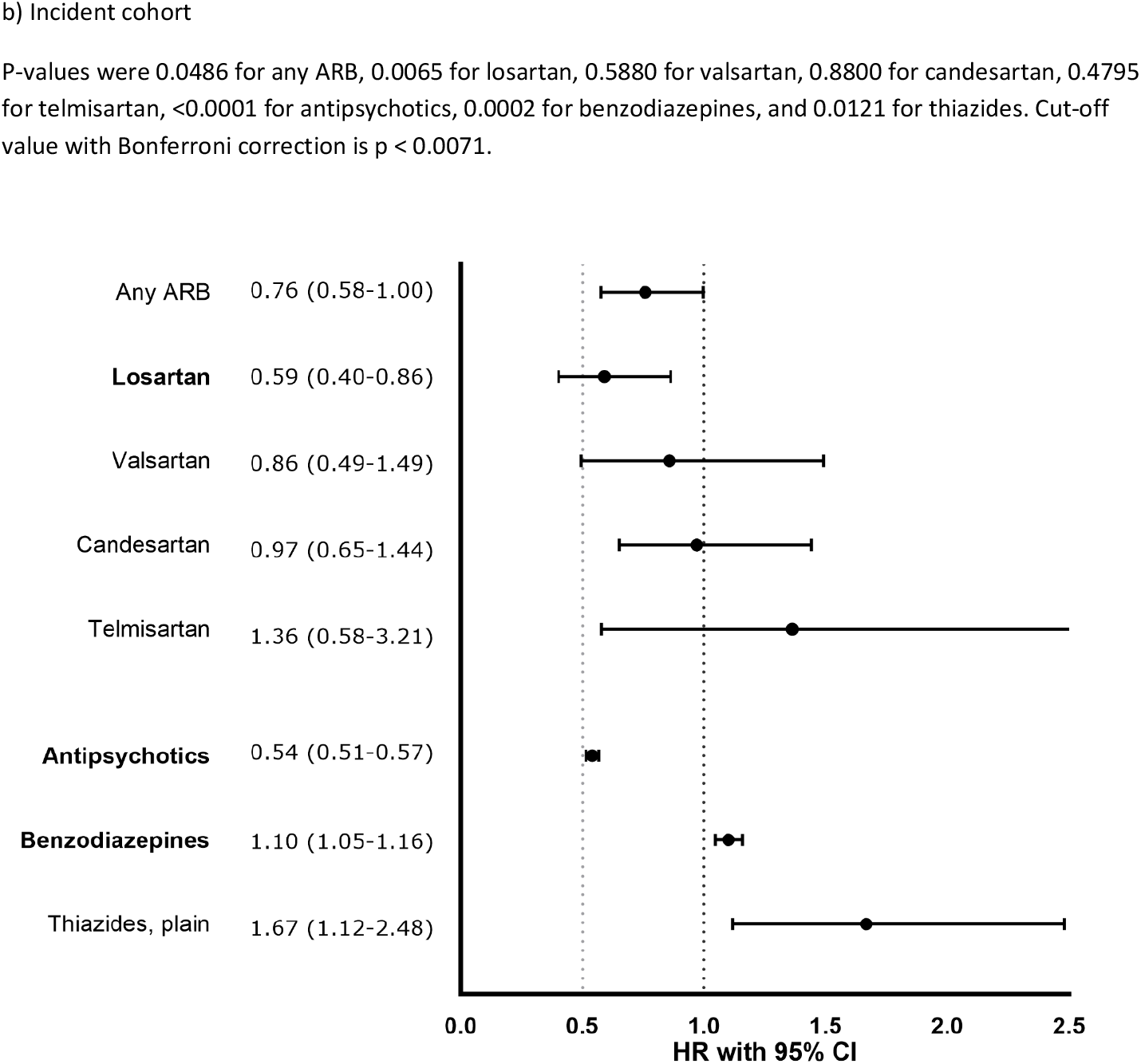
Risk of psychiatric re-hospitalization during the use of an angiotensin II receptor antagonist (ARB) or thiazide diuretic, antipsychotic, or benzodiazepine vs. no use of the drug. In bold are depicted the agents that are significant after Bonferroni correction.

## Discussion

To our knowledge, this is the first study to analyze schizophrenia-specific molecular pathways from two independent datasets, showing a consistent overlap of altered gene expression. While glutamate receptor signaling was highly abnormal only among Finnish patients compared with unrelated healthy controls in the canonical pathway analysis, GP 6 and hepatic fibrosis/hepatic stellate cell activation were the most significant pathways in the comparison between patients and healthy controls in the US dataset, and in the comparison between affected and unaffected monozygotic twins in the Finnish dataset. The finding on GP 6 pathway is in line with a large body of literature showing abnormalities in extracellular matrix (ECM) and perineuronal nets (PNNs) in schizophrenia [12,13]. ECM regulates cell migration, differentiation, neuronal plasticity, and neurite outgrowth, and a disintegrin and matrix metalloproteases with thrombospondin motif (ADAMTS) cleave ECM molecules for dynamic functional adaptations [14]. PNNs are assemblies of extracellular glycoproteins which ensheath parvalbumin-expressing GABA interneurons, and they protect neurons from oxidative stress [13]. *ADCYAP1, ADAMTS*, and *CACNA* were involved in the majority of the statistically most significant causal pathways in both Finnish and US datasets. Although both dopaminergic and GABAergic genes were expressed in both datasets, dopamine-related genes *(DRD2, COMT)* contributed to the most significant pathways only in the Finnish dataset, and GABAergic gene *(GABRA2)* only in the US dataset.

The strongest genetic association with schizophrenia has been localized in the major histocompatibility complex (MHC) locus [3], and later it was shown that this association is partially attributable to diverse alleles of the complement component (C4) genes [15]. A recent study demonstrated that C4 variants are associated with inflammation and excessive synaptic pruning mediated by monocytes differentiated to microglia direction in the iPSC-derived neurons of patients with schizophrenia [16]. In our analyses, several pathways differentiating patients from controls included pathways related to inflammation and oxidative stress. In the comparison between Finnish monozygotic twins discordant for schizophrenia, the most significant major regulator was complement C3b/4b receptor 1-like gene (CR1L). So far, CR1L has been linked to Alzheimer’s disease [17], but our results suggest that CR1L-regulated C4 activation may also have an important contribution to the development of schizophrenia. In the US dataset, cytosolic phospholipase 2 (Cpla2) was the master regulator for a causal pathway, including *ADAMTS2* and *GABRA2.* Cpla2 is involved in inflammatory responses, and it is increased in schizophrenia and autism [18]. The P2RY11-regulated pathway in the Finnish dataset and the astaxanthin-regulated pathway in the US dataset imply an abnormal purine-related molecular cascade underlying, which is in line with data from small RCTs suggesting beneficial effects for purinergic agents [19].

The results on the losartan-regulated causal pathway involving *ADCYAP1, ADAMTS9, CAGNA2D1, COMT, GRIK2, NMDAR2B*, and *VGLUT2* are in line with previous studies indicating that angiotensin II receptor antagonists may have a beneficial effect on mental disorders [20,22]. This implies that losartan and other angiotensin II receptor antagonists (ARBs) might be effective also in schizophrenia. Furthermore, our pharmacoepidemiological data showed that in the total nationwide cohort, the use of losartan and another widely used angiotensin II receptor antagonists was associated with about a 25% to 30% lower risk of hospitalization due to psychosis compared to those time periods when the same individual did not use an angiotensin II receptor antagonist. This beneficial association was even stronger in the younger first-episode cohort, showing a 41% lower risk of rehospitalization during the use of losartan. These findings were not explained by the overall temporal variation in treatment compliance since no effect was observed for thiazide diuretics used for the same indication, hypertonia. Concerning specific ARBs, the outcome was not associated with the permeability through the blood-brain-barrier (BBB). Losartan’s access through the BBB is considered rather poor in the healthy brain without trauma or neuropsychiatric illness. However, data from animal studies show that it can penetrate brain after oral administration. It inhibits epileptic seizures by preventing astrocyte activation [23], inhibiting albumin-induced PNN degradation around parvalbumin interneurons [24], and reducing BBB dysfunction [25]. It also enhances the extinction of fear memory when administered peripherally [26]. ARBs decrease brain inflammation and glutamate excitotoxicity via central and peripheral mechanisms and repurposing them for treatment of neuropsychiatric disorder may be of major immediate and translational value [27]. While losartan was associated with decreased risk of relapse in schizophrenia, especially among first-episode patients, hypothetically, its most beneficial effect should be seen in primary prevention if it could normalize the pathophysiological cascade related to down-regulated *ADCYAP1, ADAMTS*, and *CACNA* genes.

Although the number of participants was smaller in the Finnish dataset (n = 15) than in the US dataset (n = 22), the findings were substantially stronger in the Finnish dataset. This was probably due to the more homogeneous study population, resulting in less illness-irrelevant genetic noise. In our previous study, we observed that the number of sex-specific genes increased as a function of time during the maturation of cells [8]. Also, the analyses of the present study showed that the number of DEGs is higher when the neurons have matured for a longer time. Although the US dataset included several non-Caucasian controls, the results were surprisingly similar to those from the Finnish dataset. The most consistent finding across the Finnish and US datasets was the involvement of the *ADCYAP1* gene in the majority of the most significant causal pathways. *ADCYAP1* codes the pituitary adenylate cyclase-activating polypeptide (PACAP) protein, which upregulates *DISC1* (disrupted in schizophrenia 1 [28]). It also modulates dendritic spine maturation and morphogenesis [29] and has been studied extensively in schizophrenia and other psychiatric disorders [28]. One study has shown that metabotropic glutamate 2/3 agonist MGS0028 improves impairments in the novel object recognition test in mice lacking PACAP in an experimental *Adcyap1^-/-^* mouse model for schizophrenia [30]. Another study showed that PACAP-deficient mice have psychomotor and cognitive deficits and that 5-HT7 antagonist SB-269970 ameliorated these deficits [31]. This suggests that MGS0028 and 5-HT7 antagonists could be potential treatments in primary prevention before the full manifestation of illness involving dopaminergic abnormalities.

A major limitation of hiPSC-studies is the small number of subjects due to the laborious method. Our study included a total of 37 individuals, which is a rather small sample but, to our knowledge, the largest this far in schizophrenia research using hiPSC-derived neurons. Despite the sample size, findings on *ADCYAP1, ADAMTS*, and *CACNA* –related signaling pathways were surprisingly consistent in two independent datasets.

In conclusion, our results showed that while glutamatergic alterations in the Finnish dataset and GABAergic alterations in the US dataset were linked with schizophrenia in hiPSC derived neurons corresponding to cells maturing during the second trimester, no consistent signal was observed for dopamine-specific genes in both datasets. This suggests that the inhibitory-excitatory balance between glutamate and GABA may be a primary pathophysiology among patients with Caucasian origin, and dopaminergic deficits emerge later in the cascade when the illness is fully manifested. Therefore, secondary prevention with dopaminergic drugs may be too late for a full recovery, but ADCYAP1/PACAP-deficit alleviators, such as metabotropic glutamate 2/3 agonists, 5-HT7 antagonists, and angiotensin II receptor blockers might be beneficial in the prodromal phase of schizophrenia, and they should be studied in RCTs.

## Supporting information

Main Supplement

## ARTICLE INFORMATION

### Data availability

All data needed to evaluate the conclusions in the manuscript are provided in the manuscript or the supplementary material.

## Acknowledgements

We thank Laila Kaskela, Eila Korhonen, and Sara Wojciechowski for technical help in generation and characterization of the stem cell lines, and Aija Räsänen for secretarial assistance. No compensation was received outside the usual salary.

## Funding/Support

The study was partially funded by the Ministry of Social Affairs and Health, Finland, through the developmental fund for Niuvanniemi Hospital, Business Finland, Sigrid Juselius Foundation, the University of Helsinki, and the University of Eastern Finland.

## Author Contributions

JT conceived the study. KT carried out the pathway analyses. ŠL planned and supervised iPSC lines characterizations, differentiation of the neurons, and sample preparation for RNA and protein sequencing. TC, JL, ST, and JS gathered the data on twin pairs. IO and OV performed skin biopsies and rating of symptoms, MK differentiated the neurons and prepared RNA samples for sequencing. ML contributed to the interpretation of the results. AT and HT were in charge of the pharmacoepidemiological analyses. JT wrote the first draft of the manuscript. All authors contributed to the critical revision of the manuscript.

## Conflict of Interest Disclosures

The authors declare no competing interests.

## Additional information

Supplementary Online Content accompanies this paper.

## Notes

### Competing Interest Statement

The authors have declared no competing interest.

