## Supplementary material for "Molecular signaling pathways underlying schizophrenia": Main Supplement

**Supplementary Methods.** Pharmacoepidemiological analysis

**Supplementary Table 1.** Characteristics of the pharmacoepidemiological study cohort.

**Supplementary Table 2.** Definition of medications in the pharmacoepidemiological study.

**Supplementary Table 3.** Covariates used for adjusting the models.

**Supplementary Table 4.** IPA Canonical pathways and Causal networks. Tiihonen et al. [7] – Affected Twins vs. Unrelated Healthy Controls.

**Supplementary Table 5.** IPA Canonical pathways and Causal networks. Novel Finnish dataset – Affected vs. Unaffected Twins.

**Supplementary Table 6.** IPA Canonical pathways and Causal networks. Hoffman et al. [8] – Patients with Schizophrenia vs. Healthy Controls

**Supplementary Table 7a.** The genes included in the top 10 causal pathways in the Finnish dataset [7].

**Supplementary Table 7b.** The genes included in the top 10 causal pathways in the US dataset [8].

**Supplementary Table 8.** IPA Canonical pathways and Causal networks. Tiihonen et al. [7] – Unaffected Twins vs. Unrelated Healthy Controls.

**Supplementary Table 9a.** Events and incidence rates of psychiatric re-hospitalizations during drug use in the prevalent cohort.

**Supplementary Table 9b.** Events and incidence rates of psychiatric re-hospitalizations during drug use in the incident cohort.

**Supplementary Figure 1.** The most significant IPA Causal Network of ref. [7], with master regulator P2RY11, predicted participating regulators and affected genes in the data set.

**Supplementary Figure 2.** The second most significant IPA Causal Network of ref. [7], with master regulator losartan potassium, predicted participating regulators and affected genes in the data set.

**Supplementary Figure 3.** The most significant IPA Causal Network in ref. [8], with master regulator CPLA2, predicted participating regulators and affected genes in the data set.

**Supplementary Figure 4.** The second most significant IPA Causal Network of ref. [8], with master regulator alpha-adrenergic receptor, predicted participating regulators and affected genes in the data set.

**Supplementary Figure 5.** The mechanistic network of the most significant IPA upstream regulator in comparison of affected and unaffected twins, CR1L, with predicted participating regulators and affected genes in the data set.

**Supplementary Methods**

**Pharmacoepidemiological analysis**

Study population

In Finland, all residents have a personal identification code that allows linkage between several nationwide registers. The study cohort consisted of persons with schizophrenia or schizoaffective disorder treated in inpatient care between 1972 and 2014 in Finland, identified from the Hospital Discharge Register (ICD-10 diagnosis codes F20 and F25, and ICD-8 and ICD-9 diagnosis code 295* [1]. The Hospital Discharge Register is maintained by the National Institute of Health and Welfare, and it includes all inpatient hospital stays in Finland. After excluding 361 individuals without sufficient follow-up periods, the prevalent cohort included 61 889 persons (median age 46 years), and the incident cohort 8 342 persons (median age 36 years). The follow-up started for the prevalent cohort on January 1, 1996, and at first discharge from inpatient care for the incident cohort. The follow-up ended at death or on December 31, 2017, whichever occurred first. Data were analyzed from February 2 to March 18, 2020. The characteristics of the cohort are shown in Supplementary Table 1.

Exposure

The exposures were drug use periods of angiotensin II receptor blockers (ARBs), thiazide diuretics, antipsychotics and benzodiazepines. ARBs were analyzed on specific drug level, and losartan was of specific interest, because it was a master regulator of a highly significant causal pathway. To assess the putative role of general adherence to drug treatments, we used plain thiazide diuretics as a controlled exposure, since there is no evidence suggesting that thiazide diuretics have any effects on treating schizophrenia. Antipsychotics and benzodiazepines were included to assess the effectiveness of psychotropic treatments. The detailed list of drugs used is provided in Supplementary Table 2.

Information about drug use was derived from the Prescription Register, which is maintained by the Social Insurance Institution of Finland. The register includes reimbursed drug dispensations. Drug dispensations were modeled into drug use periods with PRE2DUP modeling, which takes into account daily dose in defined daily dose (DDD), dose changes, the regularity of drug dispensations, possible stockpiling of drugs and hospital stays [2]. PRE2DUP method has been validated by comparing PRE2DUP results with interview-based information on drug use.

Outcomes

The main outcome was hospitalization due to psychosis (ICD-10 codes F20-29; See Supplementary Table 3).

Covariates

In the within-individual model, individuals act as their own control and, therefore, time-invariant covariates are automatically controlled for in the study design. The model was adjusted for time-varying covariates, which included the use of antipsychotics, mood stabilizers, benzodiazepines and antidepressants, the temporal order of treatments, and time since the cohort entry date.

Statistical Analyses

In order to eliminate selection bias, we used within-individual analysis, which has been used in several previous studies [3-6]. Hospitalization due to psychosis was analyzed in a within-individual model where follow-up time was reset to zero after each outcome event. All persons who had variation in exposure and who had an outcome event during the follow-up contributed to the within-individual analyses. Within-individual models were analyzed with stratified Cox regression [3]. Hazard ratios (HR) were calculated for the hospitalization-based outcomes when the same person was using an investigated drug (angiotensin II receptor antagonist or thiazide diuretic) compared to the time periods when this same individual did not use the drug. Since it can be anticipated that the putative beneficial effect is stronger during the early phase of the illness than during a more chronic phase, we also conducted an analysis among the first-episode patients (the incident cohort).

Pertinent institutional authorities granted permissions for this research project at the Finnish National Institute for Health and Welfare (permission THL/1466/6.02.00/2013), the Social Insurance Institution of Finland (34/522/2013), and Statistics Finland (TK53-305-13).

**Supplementary Table 1.** Characteristics of the pharmacoepidemiological study cohort.

|  | Total cohort,  N (%) | Total cohort, ARB users, N (%) | Incident cohort,  N (%) | Incident cohort, ARB users, N (%) |
| --- | --- | --- | --- | --- |
| **Number of people in cohort** | 61 889 | 6865 | 8342 | 763 |
| **Age at baseline, years** |  |  |  |  |
| ≤ 35 | 17 377 (28.08%) | 1241 (18.08%) | 4043 (48.47%) | 193 (25.29%) |
| 36–55 | 27 415 (44.30%) | 3901 (56.82%) | 2534 (30.38%) | 325 (42.60%) |
| > 55 | 17 097 (27.63%) | 1723 (25.10%) | 1765 (21.16%) | 245 (32.11%) |
| **Median age (IQR), years** | 45 (34–57) | 47 (39–56) | 36 (26–52) | 48 (35–59) |
| **Male gender** | 31 104 (50.26%) | 2990 (43.55%) | 4696 (56.29%) | 355 (46.53%) |
| **Number of psychiatric hospitalizations** |  |  |  |  |
| at baseline ≤ 1 | 19 002 (30.70%) | 2244 (32.69%) | 4808 (57.64%) | 467 (61.21%) |
| 2–3 | 18 839 (30.44%) | 2078 (30.27%) | 2548 (30.98%) | 217 (28.44%) |
| > 3 | 24 048 (38.86%) | 2543 (37.04%) | 950 (11.39%) | 79 (10.35%) |
| during follow-up ≤ 1 | 32 878 (53.12%) | 3751 (54.64%) | 4557 (54.63%) | 485 (63.56%) |
| 2–3 | 9 926 (16.04) | 1046 (15.24%) | 1539 (18.45%) | 118 (15.47%) |
| > 3 | 19 085 (30.84%) | 2068 (30.12%) | 2246 (26.92%) | 160 (20.97%) |
| **Time since first SZ diagnosis, years** |  |  |  |  |
| ≤ 1 | 23 555 (38.06%) | 2348 (34.20%) | 7843 (94.02%) | 738 (96.72%) |
| 1–5 | 5 579 (9.01%) | 555 (8.08%) | 418 (5.01%) | 18 (2.36%) |
| > 5 | 32 755 (52.93%) | 3962 (57.71%) | 81 (0.97%) | 7 (0.92%) |
| **Comorbidities at baseline** |  |  |  |  |
| cardiovascular disease | 9 651 (15.59%) | 1271 (18.51%) | 1247 (14.95%) | 209 (27.39%) |
| diabetes | 3 208 (5.18%) | 451 (6.57%) | 296 (3.55%) | 57 (7.47%) |
| asthma | 1 733 (2.80%) | 199 (2.90%) | 323 (3.87%) | 34 (4.46%) |
| cancer | 1 766 (2.85%) | 170 (2.48%) | 225 (2.70%) | 33 (4.33%) |
| **Comorbidities during follow-up** |  |  |  |  |
| cardiovascular disease | 19 116 (30.89%) | 3487 (50.79%) | 1582 (18.96%) | 303 (39.71%) |
| diabetes | 8 120 (13.12%) | 1665 (24.25%) | 610 (7.31%) | 131 (17.17%) |
| asthma | 4 856 (7.85%) | 770 (11.22%) | 329 (3.94%) | 59 (7.73%) |
| cancer | 6 618 (10.69%) | 766 (11.16%) | 479 (5.74%) | 65 (8.52%) |

Abbreviations: ARB = angiotensin II receptor antagonist, SZ = schizophrenia

*Time since first SZ diagnosis indicates time since the first admission to hospital. For few patients, the first hospital treatment has lasted for several years.

**Supplementary Table 2.** Definition of medications in the pharmacoepidemiological study.

| **Drug type** | **Drug** | **ATC** |
| --- | --- | --- |
| ARB | Losartan | C09CA01, C09DA01 |
|  | Eprosartan | C09CA02, C09DA02 |
|  | Valsartan | C09CA03, C09DA03, C09DB01, C09DX01, C09DX04, |
|  | Candesartan | C09CA06, C09DA06 |
|  | Telmisartan | C09CA07, C09DA07 |
|  | Olmesartan | C09CA08, C09DB02, C09DA08 |
| Thiazide diuretics, plain | Cyclothiazide | C03AA09 |
|  | Hydrochlorothiazide | C03AA03 |
| Antidepressants | Non-selective monoamine reuptake inhibitors | N06A |
|  | Selective serotonin reuptake inhibitors |  |
|  | Monoamine oxidase inhibitors, non-selective |  |
|  | Monoamine oxidase A inhibitors |  |
|  | Other antidepressants |  |
| Antipsychotics | Phenothiazines with aliphatic side-chain | N05A |
|  | Phenothiazines with piperazine structure |  |
|  | Phenothiazines with piperidine structure |  |
|  | Butyrophenone derivatives |  |
|  | Indole derivatives |  |
|  | Thioxanthene derivatives |  |
|  | Diphenylbutylpiperidine derivatives |  |
|  | Diazepines, oxazepines, thiazepines and oxepines |  |
|  | Benzamides |  |
|  | Other antipsychotics |  |
| Mood stabilizers | Carboxamide derivatives | N03AF |
|  | Fatty acid derivatives | N03AG |
|  | Lithium | N05AN01 |
|  | Other antiepileptics | N03AX |
| Anxiolytics | Benzodiazepine derivatives | N05BA |
| Hypnotics and sedatives | Benzodiazepine derivatives | N05CD |
|  | Benzodiazepine related drugs | N05CF |

Abbreviations: ATC = Anatomical Therapeutic Chemical classification system, ARB = angiotensin II receptor antagonist.

**Supplementary Table 3.** Covariates used for adjusting the models.

| **Covariate** | **Definition** |
| --- | --- |
| Temporal order of treatments | Order of treatment continuously updated in the models, categorized as no treatment, 1st, 2nd, 3rd, >3rd |
| Concomitant use of psychotropic drugs | Antidepressants (N06A), antipsychotics (N05A excluding lithium), benzodiazepines and related drugs (N05BA, N05CD, N05CF) and mood stabilizers (valproate, carbamazepine, lamotrigine, lithium) continuously updated in the models |
| Time since cohort entry (=time since 1.1.1996 for prevalent and since the first diagnoses for incident cases | In years, categorized as ≤ 1, > 1-5, > 5, continuously updated in the models |

**Supplementary Table 7a.** The genes included in the top 10 causal pathways in the Finnish dataset [7].

| **10** |  | **9** |  | **8** |  | **7** |  | **6** |  | **5** |  | **4** |  | **3** |  | **2** |  | **1** |  |
| --- | --- | --- | --- | --- | --- | --- | --- | --- | --- | --- | --- | --- | --- | --- | --- | --- | --- | --- | --- |
| ARFGEF3  CABP7  CACNA2D1  CLMP  NEUROD6  SFRP4  SIAH3  SLC17A6  SYN2 | 0  0  3  0  2  0  0  11  12 | BHLHE22  CBWD1  COMT  GRIK2  NKX2-1  NTS | 0  0  671  11  7  19 | ADAMTS9  ADCYAP1  INSRR  NPPA  VSNL1 | 1  11  5  0  8 | HLA-A  NEUROD1  OAS3  SP100  TAC1  TRPC3 | 59  7  0  0  7  1 | CDH1  CDON  HEY2  KIT  MALAT1  MGP  NFKBIZ  NPTX1  NTN1  PPARGC1A  PRDM16 | 0  0  1  48  2  5  0  1  1  10  1 | CXCL12  DCT  GLI1  GRIN2B  GRM1  HDC  HLA-C  KCNQ3  LHX2  NTRK1  RGS4  SHH  SOX1  TFAP2A | 6  1  2  71  12  3  21  1  3  5  114  21  2  5 | CHST8  COL6A3  DRD2  LYPD1  NRN1  RASGRP1  RBP1  TFAP2C | 1  0  455  0  6  1  1  5 | ACTN2  ASPN  FNDC5  GBX2  HS3ST1  KCNQ1OT1  LHX6  MN1  MPV17L  NEGR1  NPNT  PRDM8  RAMP1  SHISA6  SOX2-OT  TBX1  TCF7L2  THSD4  ZFHX3 | 0  0  1  0  0  0  12  0  0  3  0  0  1  0  1  27  12  0  0 | EOMES  FMO5  LTBP1  NUDT19  PLAAT3  PRKG2  RGS11  RNASET2  SIX3  SLC17A7  STK17B  SV2B  SVIP  TBR1  TC2N  TSPO | 2  0  0  0  0  0  0  0  1  32  0  0  0  11  0  37 | B3GAT1  CACNG5  CDHR1  DLG2  ESRRG  FGF17  FOXP1  GPR37  ISLR  KCNJ5  LHX8  MAB21L1  ME1  MRI1  PDE7B  SFTA3  SIX6  SPINK5  TCEAL5  VWA5B1  ZNF536 | 4  2  0  8  0  0  7  1  0  0  3  2  0  0  4  0  1  0  0  0  2 |

The number above each column indicates how many times each gene is included in the top 10 pathways, and the number in the right of each gene indicates the number of publications for each gene with search term “schizophrenia” retrieved in PubMed (April 7, 2020).

**Supplementary Table 7b.** The genes included in the top 10 causal pathways in the US dataset [8].

| **9** |  | **8** |  | **7** |  | **6** |  | **5** |  | **4** |  | **3** |  | **2** |  | **1** |  |
| --- | --- | --- | --- | --- | --- | --- | --- | --- | --- | --- | --- | --- | --- | --- | --- | --- | --- |
| CACNA1A  CFLAR  CTSO  FLNC  FZD8  POSTN | 5  0  0  1  0  0 | ADAMTS2  COL12A1  COL19A1  DDIT4L  GCNT1  LHX2  MYH14  OTX1  PRKD1  RBFOX1  TMOD1 | 2  0  1  0  0  3  0  2  0  8  1 | ADCYAP1  EMILIN2  INPP5D  ITIH5  LAMA2  MME  NEUROG1  NPNT  POU4F1  QPCT  TNFRSF17 | 11  0  0  1  1  3  2  0  0  2  0 | CAT  HEY2  HFE  KCNQ5  NPAS2  STRA6 | 271  1  1  3  5  0 | ADRA1A  DSP  PAMR1  PCP4  SLC5A12  SP8  TACR3 | 14  34  0  0  0  2  1 | BST2  DSC2  NPPC  RXRG  SULF1  SV2B | 0  1  0  0  0  0 | GABRA2  PCDHGA7  SGCD  SLITRK2  SLITRK4 | 16  0  0  4  2 | KIT  MPPED2  NPIPA8 (includes others) | 48  0  0 | CPQ  NLRP2 | 2  0 |

The number above each column indicates how many times each gene is included in the top 10 pathways, and the number in the right of each gene indicates the number of publications for each gene with search term “schizophrenia” retrieved in PubMed (April 7, 2020).

**Supplementary Table 9a.** Events and incidence rates of psychiatric re-hospitalizations during drug use in the prevalent cohort.

| **Medication** | **Use time in person-years** | **Events** | **Events/10 person-years (95% CI)** |
| --- | --- | --- | --- |
| Non-use of ARB | 773 313 | 170 206 | 2.25 (2.25-2.26) |
| Any ARB | 31 307 | 3590 | 1.15 (1.13-1.16) |
| Losartan | 13453 | 1579 | 1.17 (1.16-1.19) |
| Eprosartan | 431 | 41 | 0.95 (0.86-1.04) |
| Valsartan | 5244 | 536 | 1.02 (0.99-1.05) |
| Candesartan | 8811 | 1026 | 1.16 (1.14-1.19) |
| Telmisartan | 2716 | 329 | 1.21 (1.17-1.25) |
| Olmesartan | 653 | 79 | 1.21 (1.13-1.29) |
| Poly use of ARBs | 1694 | 272 | 1.61 (1.55-1.67) |
| **Thiazide diuretics** | 3713 | 557 | 1.50 (1.46-1.54) |
| Non-use of thiazide diuretics | 820416 | 175667 | 2.14 (2.14-2.14) |
| **Antipsychotics** | 632945 | 146015 | 2.31 (2.30-2.31) |
| Non-use of antipsychotics | 191185 | 30209 | 1.58 (1.57-1.59) |
| **Benzodiazepines** | 218933 | 61993 | 2.83 (2.82-2.84) |
| Non-use of benzodiazepines | 605197 | 114231 | 1.89 (1.88-1.89) |

Abbreviation: ARB = angiotensin II receptor antagonist.

**Supplementary Table 9b.** Events and incidence rates of psychiatric re-hospitalizations during drug use in the incident cohort.

| **Medication** | **Person-years** | **Events** | **Events/10 person-years (95% CI)** |
| --- | --- | --- | --- |
| Non-use of ARB | 81087 | 20385 | 2.51 (2.50-2.52) |
| Any ARB | 2882 | 250 | 0.87 (0.83-0.90) |
| Losartan | 1291 | 92 | 0.71 (0.67-0.76) |
| Eprosartan | 31 | 0 | NA |
| Valsartan | 549 | 44 | 0.80 (0.73-0.88) |
| Candesartan | 741 | 93 | 1.26 (1.17-1.34) |
| Telmisartan | 225 | 21 | 0.94 (0.81-1.06) |
| Olmesartan | 46 | 0 | NA |
| Poly use of ARBs | 114 | 10 | 0.88 (0.71-1.05) |
| **Thiazide diuretics** | 298 | 61 | 2.05 (1.88-2.21) |
| Non-use of thiazide diuretics | 88775 | 21197 | 2.39 (2.38-2.40) |
| **Antipsychotics** | 61761 | 14633 | 2.37 (2.36-2.38) |
| Non-use of antipsychotics | 27312 | 6625 | 2.43 (2.41-2.44) |
| **Benzodiazepines** | 16148 | 4795 | 2.97 (2.94-3.00) |
| Non-use of benzodiazepines | 72925 | 16463 | 2.26 (2.25-2.27) |

**Supplementary Figure 1.** The most significant IPA Causal Network from the Finnish dataset [7], with master regulator P2RY11, predicted participating regulators and affected genes in the data set. Red and green node colors in the figure indicate increased and decreased expression in schizophrenia samples compared to control, respectively. Orange and blue color indicate predicted activation or inhibition of the regulator node and edge, respectively, whereas the yellow edge denotes inconsistent findings with the downstream target. Solid edge depicts direct and dash line indirect relationships in the IPA database.

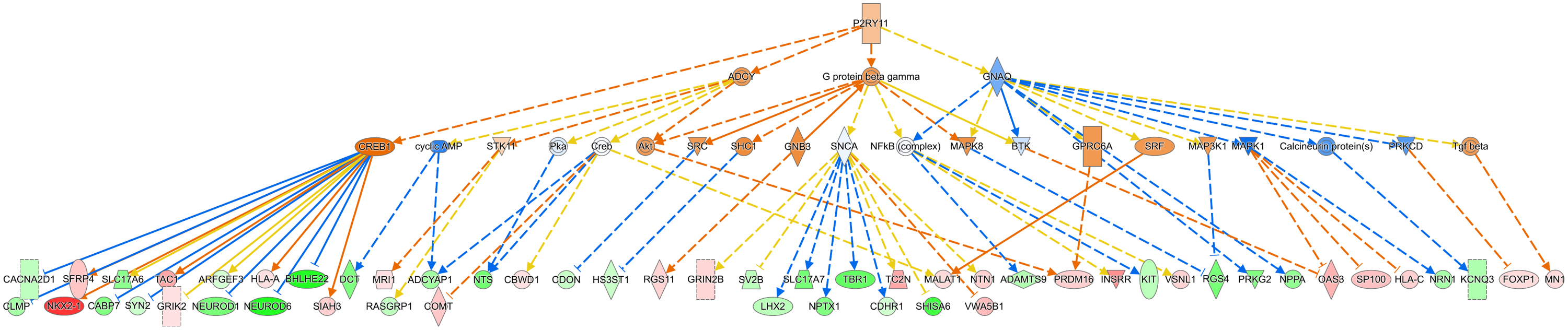

**Supplementary Figure 2.** The second most significant IPA Causal Network from the Finnish dataset [7] with master regulator losartan potassium, predicted participating regulators and affected genes in the data set. Red and green node colors in the figure indicate increased and decreased expression in schizophrenia samples compared to control, respectively. Orange and blue color indicate predicted activation or inhibition of the regulator node and edge, respectively, whereas the yellow edge denotes inconsistent findings with the downstream target. Solid edge depicts direct and dash line indirect relationships in the IPA database.

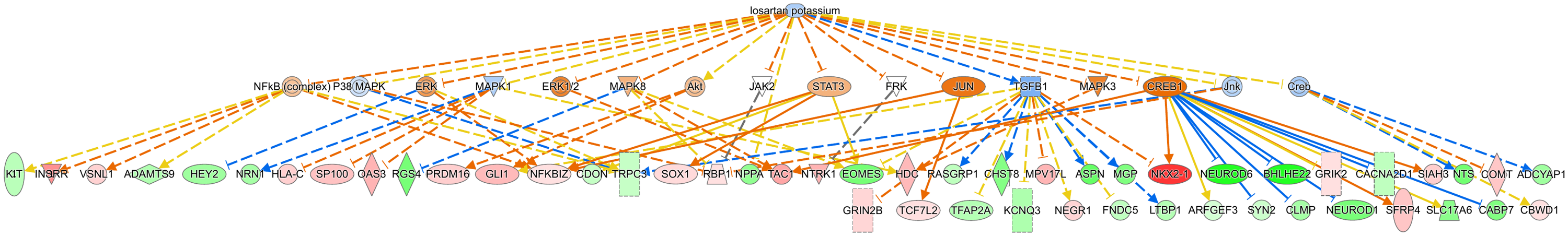

**Supplementary Figure 3.** The most significant IPA Causal Network from the US dataset [8], with master regulator CPLA2, predicted participating regulators and affected genes in the data set. Red and green node colors in the figure indicate increased and decreased expression in schizophrenia samples compared to controls, respectively. Orange and blue colors indicate predicted activation or inhibition of the regulator node and edge, respectively, whereas the yellow edge denotes an inconsistent finding with the downstream target. Solid edges depict direct and dash lines indirect relationships in the IPA database.

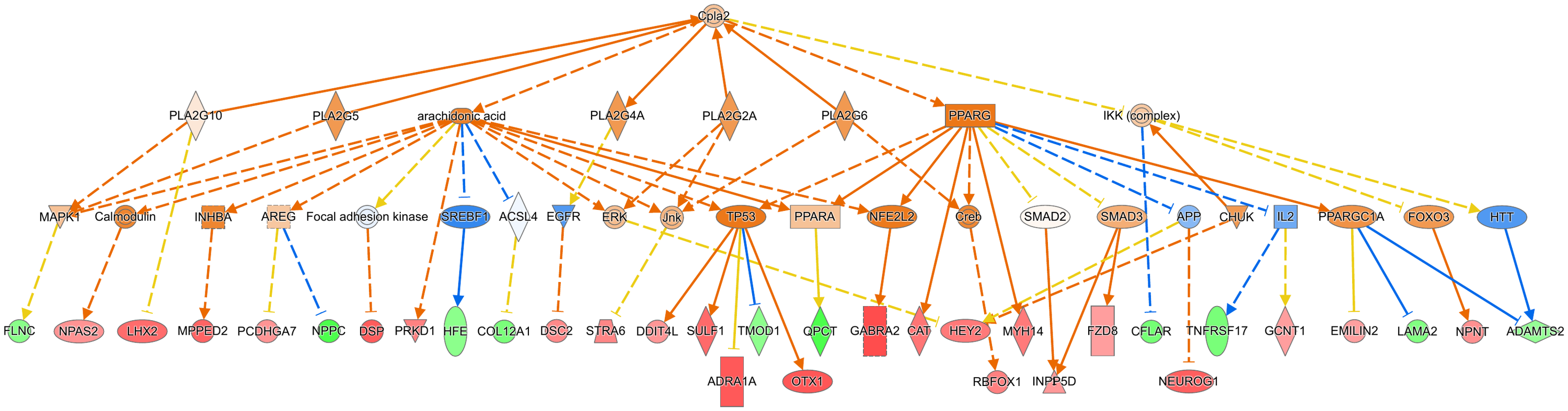

**Supplementary Figure 4.** The second most significant IPA Causal Network of from the US dataset [8], with master regulator alpha-adrenergic receptor, predicted participating regulators and affected genes in the data set. Red and green node colors in the figure indicate increased and decreased expression in schizophrenia samples compared to controls, respectively. Orange and blue colors indicate predicted activation or inhibition of the regulator node and edge, respectively, whereas the yellow edge denotes an inconsistent finding with the downstream target. Solid edges depict direct and dash lines indirect relationships in the IPA database.

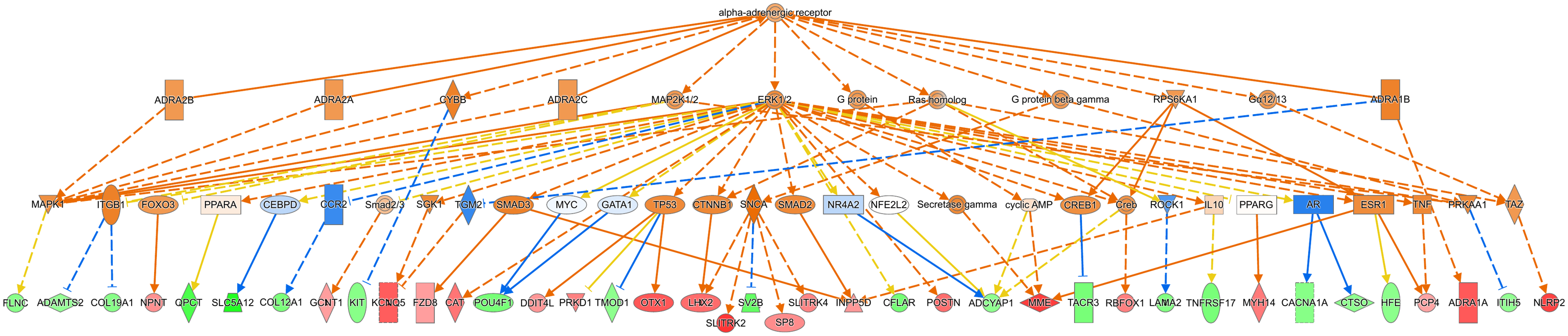

**Supplementary Figure 5.** The mechanistic network of the most significant IPA upstream regulator in comparison of affected and unaffected twins, CR1L, with predicted participating regulators and affected genes in the data set. Red and green node colors in the figure indicate increased and decreased expression in affected twins compared to unaffected twins, respectively. Orange and blue color indicates predicted activation or inhibition of the regulator node and edge, respectively, whereas yellow edge denotes inconsistent finding with the downstream target. Solid edge depicts direct and dash line indirect relationships in the IPA database.

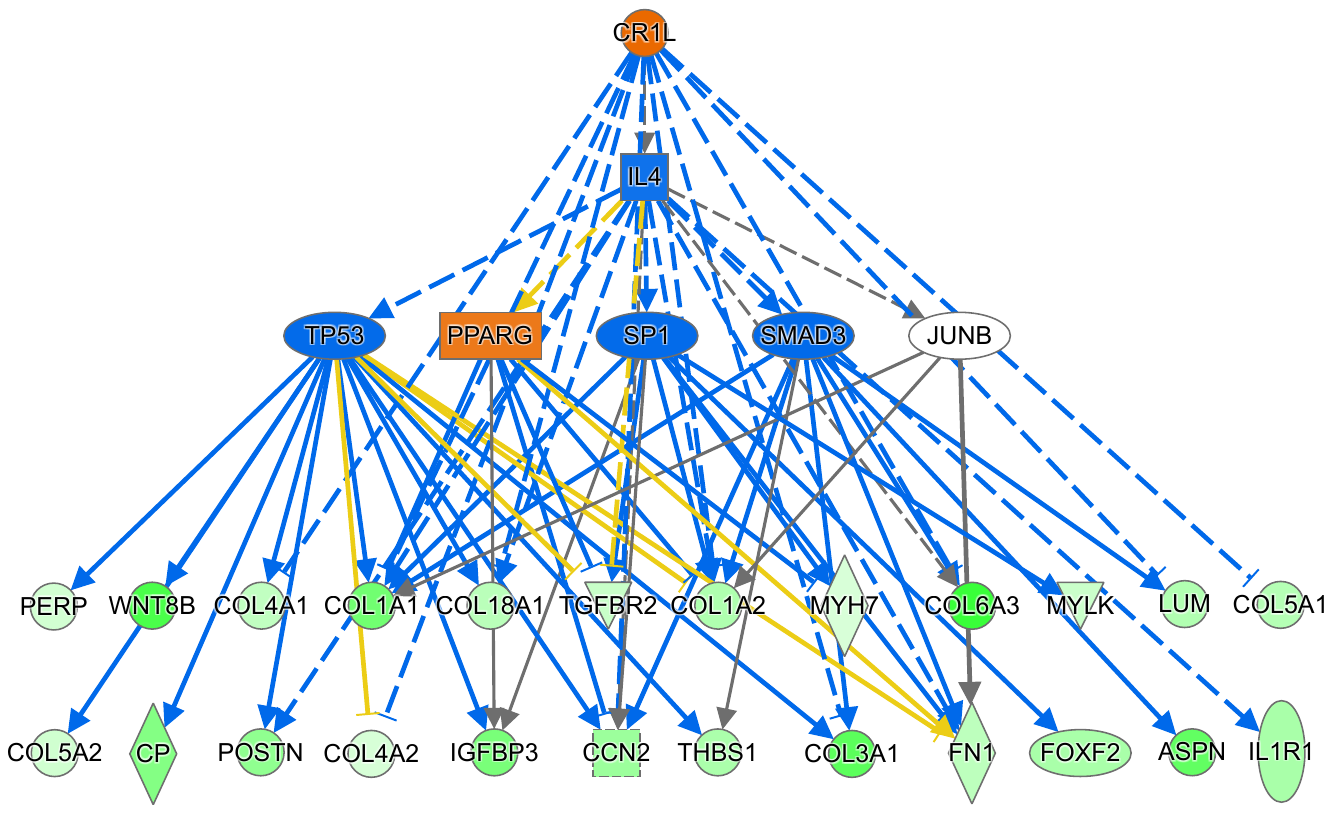
